# FLD Is Required for N-Hydroxypipecolic Acid Accumulation and Associated Growth Defects in Arabidopsis *pmr4* Mutant

**DOI:** 10.64898/2026.07.18.739330

**Authors:** Baofang Fan, Zhixiang Chen

## Abstract

The Arabidopsis mutants lacking the pathogen-inducible callose synthase POWDERY MILDEW RESISTANCE 4 (PMR4)/GLUCAN SYNTHASE-LIKE 5 (GSL5) exhibit enhanced disease resistance accompanied by reduced growth, spontaneous necrosis, and premature senescence. This autoimmune phenotype of *pmr4* is associated with a high accumulation of N-hydroxypipecolic acid (NHP) and a moderate increase in salicylic acid. We previously reported three suppressors of *pmr4* (*spm*), *PAD4* and the NHP biosynthetic genes *ALD1* and *FMO1*, whose loss-of-function mutations abolish NHP accumulation and completely suppress the growth defects of *pmr4*, demonstrating that NHP is critical to the autoimmune growth phenotype. Here, we characterize a fourth suppressor, *spm4*, which similarly fully suppresses the growth defects of *pmr4*. Map-based cloning and molecular genetic analyses revealed that *spm4* carries a mutation in *FLOWERING LOCUS D* (*FLD*). Reintroduction of a wild-type FLD allele into the *pmr4 spm4* background restored both NHP accumulation and the characteristic growth defects of *pmr4*. Conversely, independent *FLD* loss-of-function alleles generated by CRISPR/Cas-mediated genome editing suppressed NHP accumulation and rescued the autoimmune growth defects of *pmr4*, confirming that FLD is required for manifestation of the *pmr4* phenotype. Beyond its established role in flowering-time regulation, *FLD* has been implicated in plant immunity, particularly in systemic acquired resistance (SAR), although the mechanistic basis of its immune function remains poorly understood. Our findings identify FLD as an essential positive regulator of NHP accumulation, thereby providing a mechanistic link between FLD-dependent epigenetic regulation, NHP-mediated signaling, and SAR.

## INTRODUCTION

Plants rely on a two-tiered innate immune system in defense against microbial pathogens. First, plants recognize pathogen-associated molecular patterns, such as flagellin or chitin, through cell-surface pattern recognition receptors to activate pattern-triggered immunity (PTI)(Jones et al., 2024). In response, pathogens deliver effectors into plant cells to suppress PTI. Some of these effectors are recognized by intracellular nucleotide-binding leucine-rich repeat receptors (NLRs) to activate effector-triggered immunity (ETI), which is often characterized by localized programmed cell death known as the hypersensitive response(Jones et al., 2024). Activation of PTI and ETI induces immune responses, including increased biosynthesis of salicylic acid (SA), not only at the infection sites but also in distal uninfected tissues to establish systemic acquired resistance (SAR)(Hartmann and Zeier, 2019;Huang et al., 2020;Shields et al., 2022). As a central regulator of SAR, SA functions within an interconnected signaling network, acting synergistically with other defense signal molecules, including *N*-hydroxypipecolic acid (NHP) (Hartmann and Zeier, 2019;Zeier, 2021;Spoel and Dong, 2024).

Activation of plant immune responses is often associated with reduction in growth and reproductive fitness, a phenomenon commonly referred to as the growth–defense trade-offs (Huot et al., 2014;Karasov et al., 2017;He et al., 2022;Shields et al., 2022). Consequently, immune signaling pathways are tightly regulated and maintained at low basal levels in the absence of pathogen challenge (van Wersch et al., 2016;Freh et al., 2022). Multiple regulatory mechanisms, including transcriptional, post-transcriptional, and epigenetic controls, act to prevent inappropriate activation of defense responses. Disruption of these regulatory processes or gain-of-function mutations in immune receptors can result in constitutive activation of defense signaling, leading to a class of mutants collectively known as autoimmune mutants(van Wersch et al., 2016;Freh et al., 2022). These mutants typically exhibit enhanced disease resistance accompanied by elevated expression of defense-related genes, increased accumulation of immune signaling molecules such as SA and NHP, spontaneous cell death, and severe growth defects. Well-characterized examples include the NLR gain-of-function mutants *snc1* (*suppressor of npr1-1, constitutive 1*) (Li et al., 2001;Li et al., 2007;Zhu et al., 2010), *ssi4* (suppressor of salicylic acid insensitivity 4) (Zhou et al., 2004;Zhou et al., 2008), and *cpr5* (*constitutive expression of pr genes 5*) (Bowling et al., 1997;Clarke et al., 2001;Yoshida et al., 2002), all of which display constitutive defense activation and dwarf phenotypes. Similarly, mutations in negative regulators of immunity, such as *bon1* (*bonzai1*) (Yang and Hua, 2004;Lee and McNellis, 2009;Liu et al., 2025) and *acd6* (*accelerated cell death 6*) (Rate et al., 1999;Lu et al., 2003;Jasinski et al., 2020;Lyu et al., 2024), also result in autoimmune responses associated with enhanced resistance and stunted growth. Genetic analyses of these autoimmune mutants have provided fundamental insights into the structure, activation mechanisms, and signaling functions of immune receptors, as well as the complex regulatory networks that coordinate plant growth and immunity (van Wersch et al., 2016).

One autoimmune mutant that has provided important insight into the regulation of plant immunity is *powdery mildew resistance 4* (*pmr4*), which is disrupted in the pathogen-inducible callose synthase gene *PMR4* (also known as *GLUCAN SYNTHASE-LIKE 5*, *GSL5*) (Jacobs et al., 2003;Nishimura et al., 2003). Although pathogen-induced callose deposition has long been regarded as a key component of plant defense (Luna et al., 2011;Wang et al., 2021), loss of PMR4 in Arabidopsis and other plants unexpectedly results in enhanced resistance to fungal, oomycete, bacterial, and protist pathogens (Jacobs et al., 2003;Nishimura et al., 2003;Huibers et al., 2013;Martinez et al., 2020;Santillan Martinez et al., 2020;Li et al., 2022;Wu et al., 2025). These findings suggest that PMR4 plays a conserved role in modulating immune responses beyond its direct function in cell wall reinforcement. Studies in Arabidopsis revealed that the powdery mildew resistance of *pmr4* results from enhanced SA signaling based on its suppression by mutations in *PAD4* (*PHYTOALEXIN DEFICIENT 4*), a key regulator of SA signaling, and by transgenic expression of the bacterial salicylate hydroxylase gene *nahG* (Nishimura et al., 2003).

Our interest in *PMR4* arose from studies of the endoplasmic reticulum (ER)-localized UBAC2 proteins, which function in ER protein quality control and are required for pathogen-induced callose deposition through regulation of PMR4 accumulation (Wang et al., 2019). During these studies, we observed that although *pmr4* plants were largely indistinguishable from wild-type (WT) plants during the first five weeks of growth, they subsequently developed severe growth defects characterized by stunted growth, spontaneous necrosis, and premature senescence (Fan et al., 2025). Exploiting these distinctive growth phenotypes, we conducted genetic suppressor screens and identified mutations in *PAD4* and the NHP biosynthetic genes *AGD2-LIKE DEFENSE RESPONSE PROTEIN 1* (*ALD1*) and *FLAVIN-DEPENDENT MONOOXYGENASE 1* (*FMO1*) that completely restored normal growth and development in *pmr4* (Fan et al., 2025). Interestingly, while disruption of *PAD4* abolished both the growth defects and enhanced disease resistance of *pmr4*, loss of *ALD1* or *FMO1* fully suppressed the growth defects yet retained much of the powdery mildew resistance phenotype(Fan et al., 2025). In contrast, mutation of the SA biosynthetic gene *SA INDUCTION DEFICIENT 2* (*SID2*)*/ISOCHORISMATE SYNTHASE* 1(*ICS1*) had little effect on either the growth defects or disease resistance of *pmr4* (Fan et al., 2025), despite earlier evidence implicating SA signaling in *pmr4*-mediated immunity (Nishimura et al., 2003). However, simultaneous disruption of both NHP and SA biosynthesis eliminated the enhanced disease resistance of *pmr4* (Fan et al., 2025), indicating that these two defense pathways function redundantly in resistance while contributing differently to the associated fitness costs. Consistent with this model, pipecolic acid (PIP), the immediate precursor of NHP, accumulated more than 200-fold in *pmr4* at the onset of growth defects, whereas SA levels increased only approximately 3-fold(Fan et al., 2025). Together, these findings demonstrate that massive accumulation of NHP is the primary driver of the autoimmune growth defects of *pmr4*, whereas NHP and SA make redundant contributions to disease resistance. More broadly, these studies established *pmr4* as a valuable genetic model for dissecting the mechanisms underlying growth-defense tradeoffs and for distinguishing the roles of NHP and SA in plant immunity.

In the present study, we characterized an additional suppressor, *suppressor of pmr4 4* (*spm4*), identified from our genetic screens. Like *ald1* and *fmo1*, the *spm4* mutation completely restored normal growth and development in the *pmr4* background. Through map-based cloning and molecular characterization, we identified *spm4* as a mutant allele of *FLOWERING LOCUS D* (*FLD*), a gene best known for its role in flowering-time regulation through chromatin-mediated gene silencing (Yu et al., 2011;Xu et al., 2023). In addition to its developmental functions, FLD has been implicated in plant immunity and systemic acquired resistance, although the molecular basis of its immune function remains incompletely understood (Singh et al., 2013;Singh et al., 2014b;Kumar et al., 2025). Here, we demonstrate that FLD is required for the strong accumulation of NHP in *pmr4* and for the associated autoimmune growth defects. These findings identify FLD as a critical positive regulator of NHP-mediated immune responses and provide new insight into the mechanisms linking epigenetic regulation, immune signal accumulation, and the growth–defense tradeoff in plants.

## RESULTS

### Isolation and characterization of *spm4-1*, a suppressor of *pmr4* growth defects

In our previous genetic screens for suppressors of the autoimmune phenotype of *pmr4/gsl5* mutants, we identified three recessive suppressor mutants from an EMS (ethyl methanesulfonate)-mutagenized *gsl5-1* population. Molecular characterization revealed that *spm1*, *spm2*, and *spm3* carry mutations in *PAD4*, *ALD1*, and *FMO1*, respectively(Fan et al., 2025). These suppressors established that both PAD4-dependent signaling and NHP biosynthesis are required for the development of the severe growth defects associated with *pmr4*. During the same screens, we isolated an additional suppressor mutant, designated *spm4-1*, which displayed complete suppression of the characteristic autoimmune growth phenotype of *pmr4*.

Like the previously identified *spm* mutants, *gsl5-1 spm4-1* plants exhibited normal vegetative growth and development. The severe dwarfism, spontaneous necrotic lesions, and premature senescence that characterize mature *gsl5* plants were completely absent in the *gsl5 spm4-1* double mutant (Figure 1). Instead, *gsl5 spm4-1* plants were comparable in size and overall appearance to WT plants throughout development, with no obvious abnormalities in leaf morphology or plant architecture (Figure 1). An additional phenotype was consistently observed, however, in that *gsl5 spm4-1* plants flowered substantially later than WT plants under our growth conditions (Figure 1), suggesting that the causal mutation may affect a developmental regulator in addition to suppressing the autoimmune phenotype of *pmr4/gsl5*.

**Figure 1.**
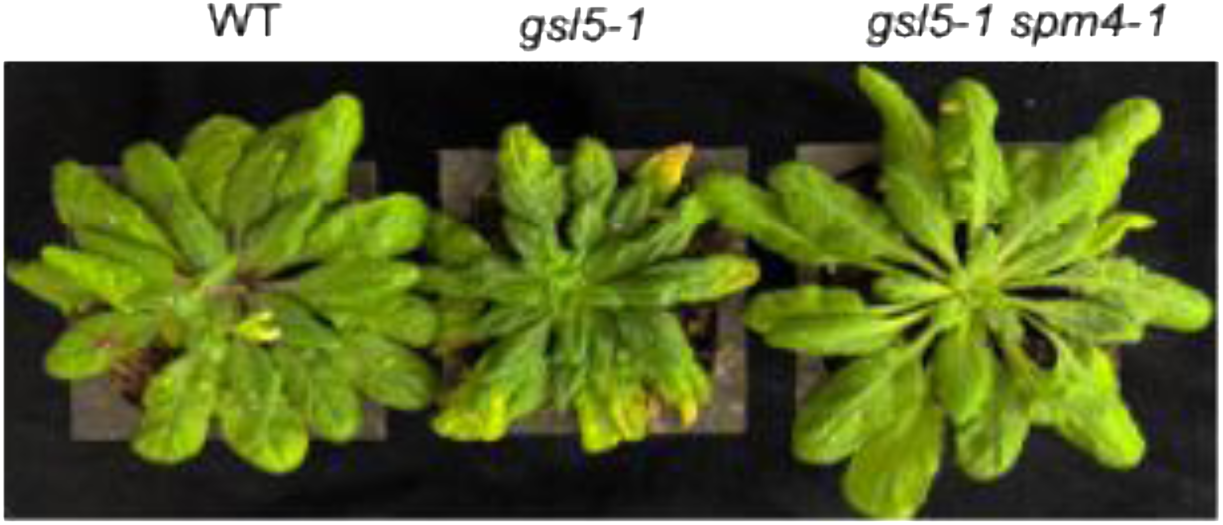
Suppression of the *gsl5-1* growth defect by the *spm4-1* suppressor mutation. Representative 9-week-old WT, *gsl5-1* and *gsl5-1 spm4-1* plants are shown (left to right). The *spm4-1* mutation suppresses the growth defects of *gsl5-1*. Notably, WT and *gsl5-1* plants had already bolted by 9 weeks of age, whereas the *gsl5-1 spm4-1* double mutant remained in the vegetative stage.

To determine the genetic basis of the suppressor phenotype, *gsl5-1 spm4-1* plants were crossed with the parental *gsl5-1* mutant. All F₁ plants displayed the characteristic *gsl5* growth defects, indicating that the suppressor mutation is recessive. Analysis of the segregating F₂ population revealed segregation patterns consistent with suppression by a single recessive nuclear mutation. These results demonstrated that the complete suppression of the *gsl5-1* growth defects in *spm4-1* is caused by a single recessive mutation, thereby facilitating subsequent map-based cloning of the affected gene.

### Mapping of *spm4-1* mutation

To identify the causal mutation responsible for the *spm4-1* suppressor phenotype, we employed bulked segregant analysis combined with next-generation sequencing (BSA-NGS). Genomic DNA samples from F_2_ individuals displaying either the original growth defects of *gsl5-1* or the normal growth of *gsl5-1 spm4-1* were pooled and subjected to whole-genome sequencing. Linkage analysis based on single nucleotide polymorphism (SNP) frequencies mapped the *spm4-1* mutation to the upper arm of chromosome III. Examination of candidate mutations within the mapped interval identified a nucleotide substitution in *FLOWERING LOCUS D* (*FLD*), a gene encoding a putative lysine-specific histone demethylase involved in epigenetic regulation. Specifically, *spm4-1* carries a G-to-A transition in the second exon of *FLD* at nucleotide position 1980 from the start codon (Figure 2A), resulting in the conversion of a tryptophan codon into a premature stop codon (Figure 2B). The predicted truncated protein terminates after amino acid residue 607 and lacks the C-terminal portion of FLD (Figure 2B), suggesting that *spm4-1* is a loss-of-function allele.

**Figure 2.**
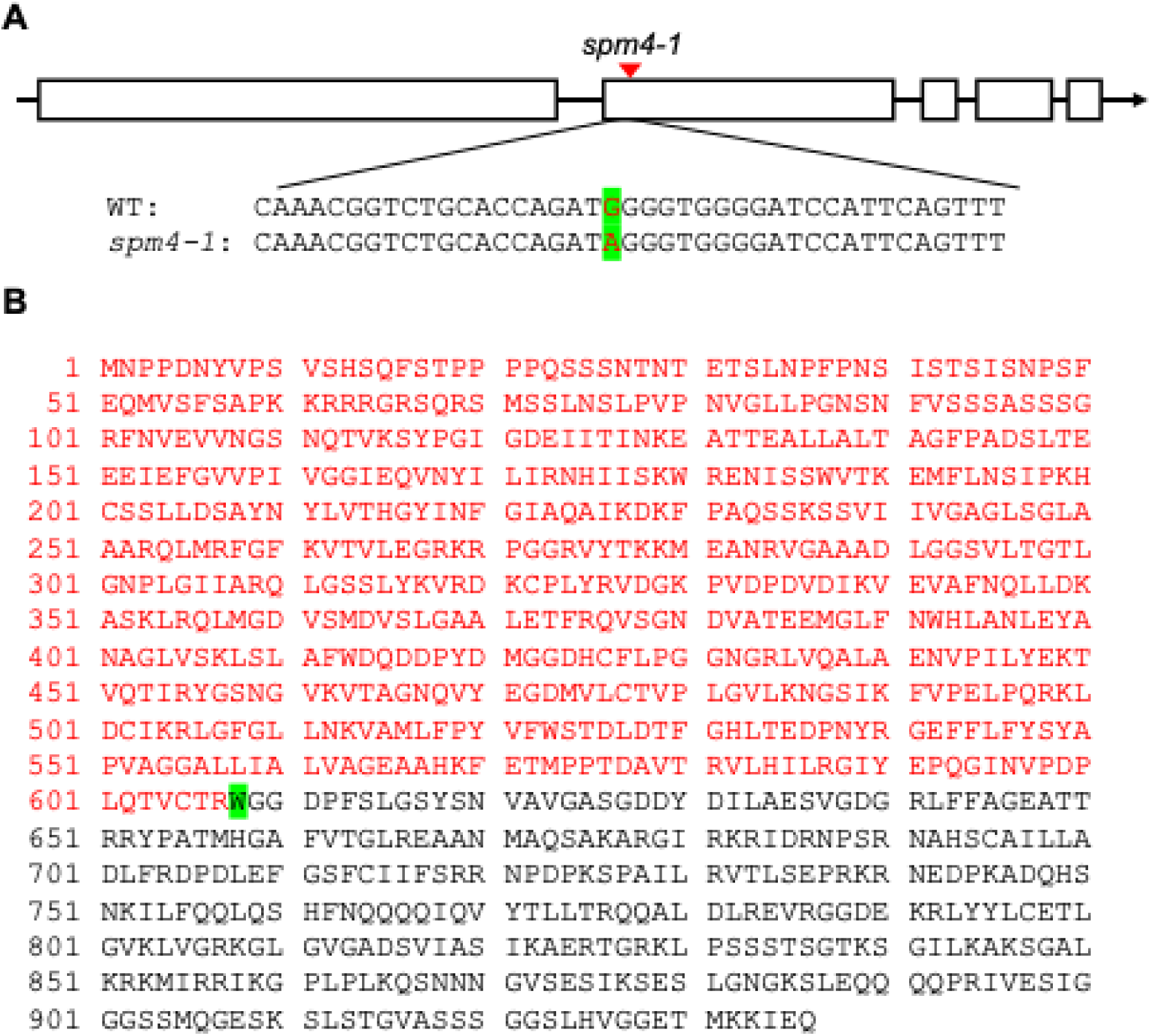
Identification of the *spm4-1* mutation. BSA-NGS mapped the *spm4-1* mutation to a genomic interval containing a G-to-A transition in the second exon of *FLD* at nucleotide position 1980 relative to the start codon **(A)**. This single-nucleotide substitution converts a tryptophan codon (TGG) into a premature stop codon (TAG), resulting in truncation of the FLD protein. A schematic of the *FLD* gene structure and the position of the *spm4-1* mutation is shown in **A.** The predicted truncated FLD protein consists of the N4erminal 607 amino acids, highlighted in red in **B**, and lacks the C-terminal portion of the wild-type protein.

### Validation of SPM4 as epigenetic regulator FLD

To determine whether disruption of *FLD* is responsible for suppression of the *pmr4/gsl5* phenotype, we performed genetic complementation by introducing a WT genomic copy of *FLD* driven by the *CaMV 35S* promoter into *gsl5-1 spm4-1* plants (Figure 3). Multiple independent transgenic lines carrying the *FLD* transgene exhibited restoration of the characteristic *gsl5-1* autoimmune phenotype. In contrast to the vigorous growth of *gsl5-1 spm4-1* plants, the complemented lines displayed severe growth reduction accompanied by spontaneous necrosis and premature senescence, closely resembling the original *gsl5-1* mutant. Reverse transcription quantitative PCR (RT-qPCR) analysis showed that, because the *FLD* transgene is driven by the constitutive *CaMV 35S* promoter, some transgenic lines accumulated *FLD* transcripts at levels substantially higher than those observed for the native *FLD* gene. Although such overexpression could potentially contribute to growth abnormalities, several independent transgenic lines expressed *FLD* at levels comparable to those of the native *FLD* mutated gene in *gsl5-1 spm4-1*(Figure 3A) yet still exhibited the typical *gsl5-1* growth phenotypes (Figure 3B). These results indicate that restoration of *FLD* function, rather than transgene overexpression, is sufficient to reestablish the *gsl5-1* phenotype and demonstrate that the *spm4-1* suppressor phenotype is caused by the mutation in *FLD*.

**Figure 3.**
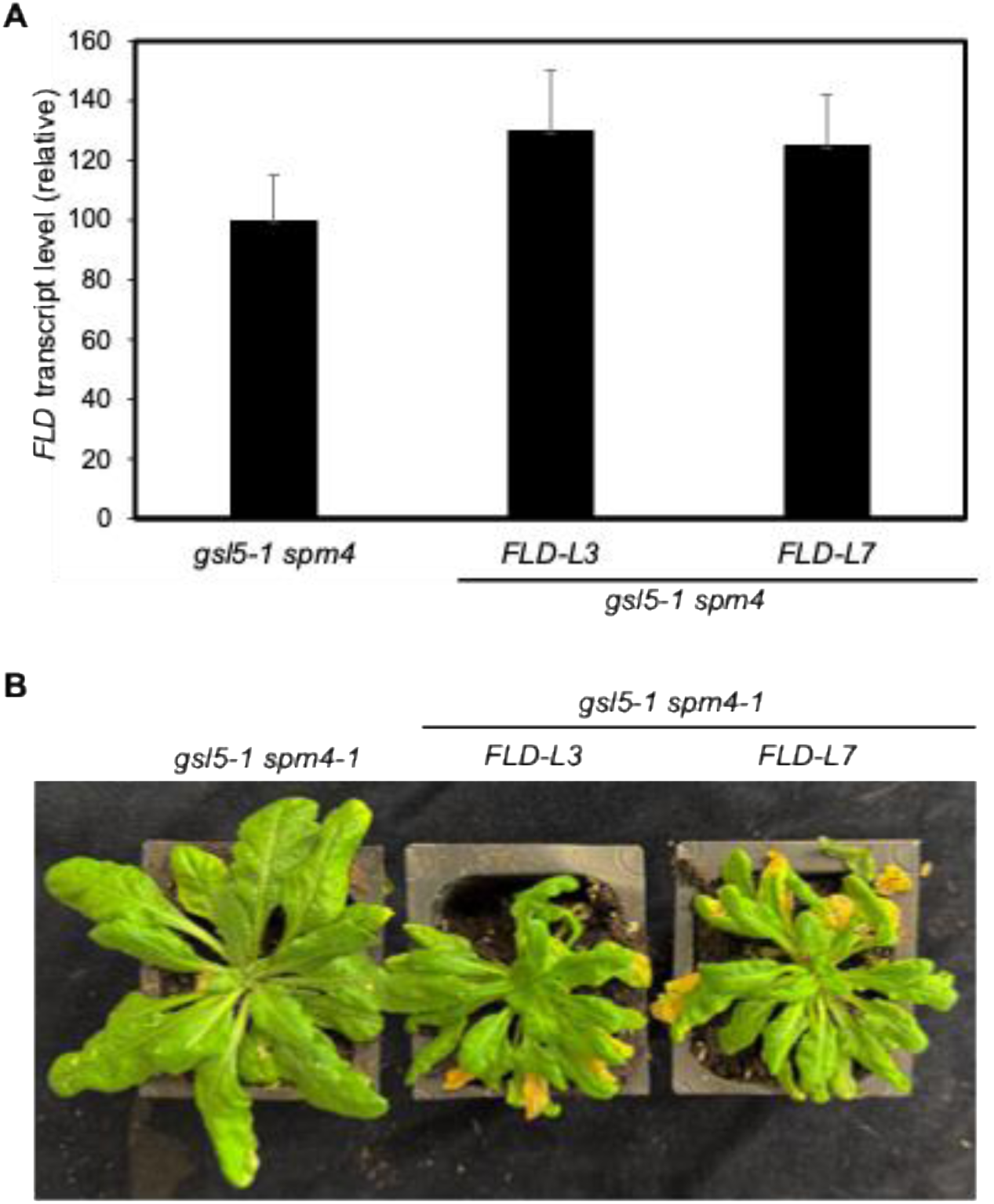
Complementation of the *gsl5-1 spm4-1* double mutant with the WT *FLD* gene restores the *gsl5-1* growth phenotype. Two independent transgenic lines, *FLD-L3* and *FLD-L7,* expressing a WT *FLD* transgene in the *gsl5-1 spm4-1* double mutant background were analyzed. RT-qPCR showed that *FLD* transcript levels in both transgenic lines were only about 30% higher than those in the *gsl5-1 spm4-1* mutant (**A**). Introduction of the functional WT *FLD* gene restored the characteristic *gsl5-1* growth defects in both *gsl5-1 spm4-1 FLD-L3* and *gsl5-1 spm4-1 FLD-L7* plants (**B**).

To further verify the identity of *SPM4*, we generated additional *FLD* mutant alleles in the *gsl5-1* background using CRISPR/Cas9-mediated genome editing (Figure 4). Two guide RNAs targeting the 5′ region of the *FLD* coding sequence within the first exon were introduced into *gsl5-1* plants. Sequence analysis of independent transformants identified two edited alleles, designated *spm4-2* and *spm4-3*, carrying a single cytosine deletion and a single adenine insertion, respectively, at nucleotide position 184 downstream of the translational start site (Figure 4A). Both mutations are predicted to cause frameshifts that result in premature termination of *FLD* translation. Consistent with the phenotype of *gsl5-1 spm4-1*, both *gsl5-1 spm4-2* and *gsl5-1 spm4-3* plants completely suppressed the autoimmune phenotype of *gsl5-1* (Figure 4B). The double mutants exhibited normal growth and lacked the dwarfism, spontaneous necrosis, and premature senescence characteristic of *gsl5-1* (Figure 4B). Together with the complementation results, these genome-editing experiments provide independent genetic evidence that *SPM4* corresponds to *FLD* and demonstrate that *FLD* function is required for the manifestation of the autoimmune growth defects associated with *gsl5-1*.

**Figure 4.**
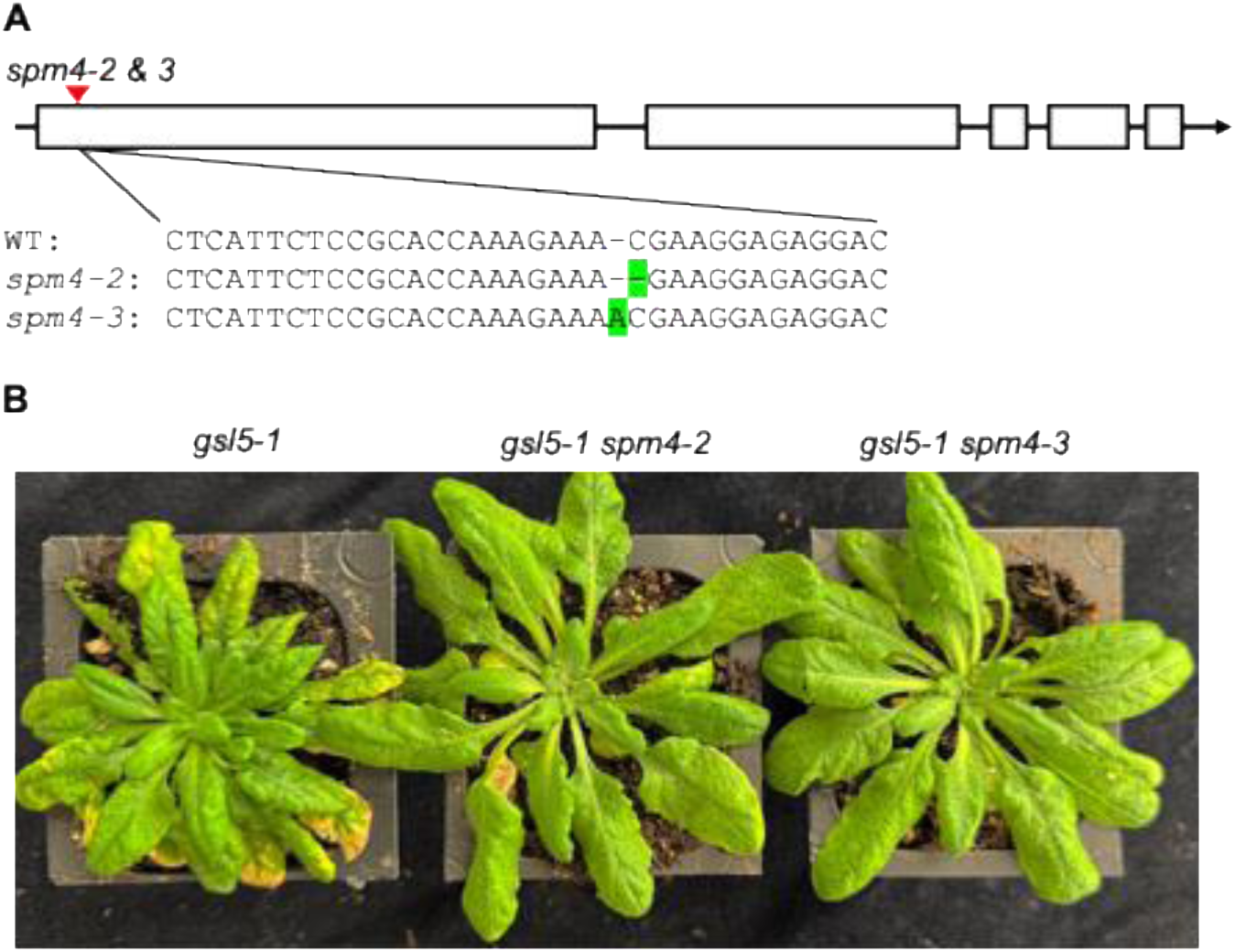
Suppression of the *gsl5-1* growth defect by the *spm4-2* and *spm4-3* suppressor mutations generated by CRISPR/cas9. The *spm4-2* and *spm4-3* mutants carry a single cytosine deletion and a single adenine insertion, respectively, at nucleotide position 184 downstream of the translational start site (**A**). Both mutations are predicted to cause frameshifts that result in premature termination of *FLD* translation. Representative 9-week-old *gsl5-1*, *gsl5-1 spm4-2,* and *gsl5-1 spm4-2* plants are shown (left to right). The *spm4-2* and *spm4-3* mutations suppress the growth defects of *gsl5-1*(**B**).

### FLD is required for the dramatic accumulation of PIP in *gsl5-1*

In our previous study, metabolic profiling revealed a striking accumulation of PIP, the direct precursor of NHP, in *pmr4/gsl5* mutants. Notably, PIP levels increased by more than 200-fold in five-week-old *gsl5* plants, before the appearance of visible growth defects, necrosis, or premature senescence (Fan et al., 2025). Genetic analyses further demonstrated that disruption of the NHP biosynthetic genes *ALD1* and *FMO1* completely suppressed the autoimmune growth phenotype of *gsl5*, establishing a critical role for the NHP pathway in the development of the mutant phenotype (Fan et al., 2025). Together, these findings indicated that the massive accumulation of NHP is the primary driver of the growth defects associated with loss of PMR4/GSL5 function (Fan et al., 2025).

Because loss of *FLD* completely suppressed the growth defects of *gsl5-1*, we next investigated whether *FLD* is also required for the elevated accumulation of NHP in the mutant. As NHP is present at lower abundance and is more difficult to quantify accurately(Chen et al., 2018;Wang et al., 2018;Holmes et al., 2021), we measured the levels of PIP, which accumulates to substantially higher levels and serves as a reliable indicator of activation of the NHP biosynthetic pathway(Chen et al., 2018;Hartmann et al., 2018;Wang et al., 2018;Brambilla et al., 2023). Consistent with our previous findings(Fan et al., 2025), five-week-old *gsl5-1* plants accumulated dramatically elevated levels of PIP, exceeding those detected in wild-type plants by more than 200-fold (Figure 5). In contrast, PIP accumulation was greatly reduced in the *gsl5-1 spm4-1* double mutant, approaching the low levels observed in wild-type plants (Figure 5). Importantly, introduction of a WT *FLD* genomic clone into the *gsl5-1 spm4-1* background restored the hyperaccumulation of PIP to levels comparable to those of *gsl5-1* (Figure 5). As expected, PIP accumulation was also strongly suppressed in the *gsl5-1 spm4-2* CRISPR mutant line (Figure 5).

**Figure 5.**
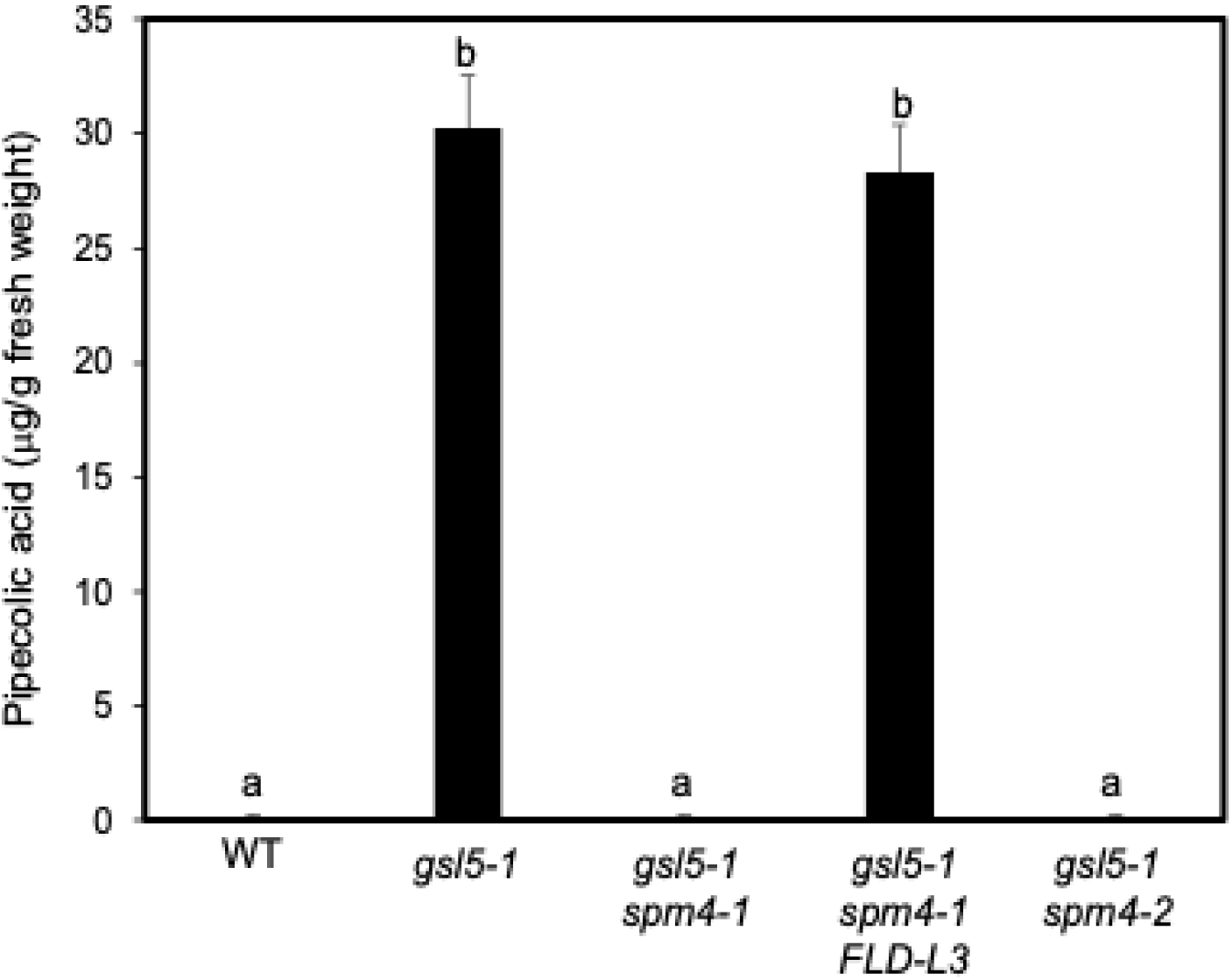
Accumulation of pipecolic acid in WT and mutants. Pipecolic acid (PIP) contents in “weeks old WT and mutant plants. Means and SE were calculated from average pipecolic acid contents determined from three samples for each genotype. According to Duncan’s multiple range test (*P* = 0.01), means of pipecolic acid contents differ significantly if they are indicated with different letters.

The close correlation between PIP accumulation and the growth phenotypes of the different genotypes strongly suggests that FLD functions upstream of NHP biosynthesis in *gsl5-1*. Loss of FLD function suppresses both the excessive accumulation of PIP and the associated autoimmune growth defects, whereas restoration of FLD activity re-establishes both traits. These results demonstrate that FLD is required for the dramatic activation of the PIP/NHP pathway in *gsl5-1* and provide a mechanistic explanation for its role in the manifestation of the autoimmune growth phenotype.

## DISCUSSION

Our previous studies established *pmr4/gsl5* as an unusual autoimmune mutant in which disruption of a pathogen-inducible callose synthase leads not only to enhanced disease resistance but also to severe growth defects associated with strong accumulation of NHP, based on the extraordinarily high levels of its immediate precursor PIP(Fan et al., 2025). Genetic analyses demonstrated that the autoimmune phenotype of *pmr4* depends on both *PAD4* and the NHP biosynthetic pathway, as mutations in *PAD4*, *ALD1*, and *FMO1* completely suppressed the stunted growth, necrosis, and premature senescence of *pmr4*(Fan et al., 2025). These studies further revealed distinct contributions of NHP and SA to the *pmr4* phenotypes. Whereas loss of NHP biosynthesis restored normal growth while retaining much of the disease resistance, simultaneous disruption of both NHP and SA biosynthesis abolished the enhanced resistance phenotype(Fan et al., 2025). Together, these findings established that massive activation of the NHP pathway is primarily responsible for the growth defects of *pmr4*, whereas NHP and SA function redundantly in disease resistance(Fan et al., 2025). In the present study, the identification of *SPM4* as *FLD* adds a new component to this regulatory pathway and demonstrates that FLD is required for the dramatic accumulation of PIP/NHP and the associated autoimmune growth defects of *pmr4*. The complete suppression of both growth defects and PIP accumulation by multiple independent *fld* alleles (Figures 1 & 4), together with restoration of both phenotypes by a wild-type *FLD* transgene (Figure 3), firmly places FLD upstream of NHP overaccumulation in *pmr4*.

FLD was originally identified as a key regulator of flowering time and belongs to a family of proteins related to mammalian lysine-specific histone demethylases (He et al., 2003;He et al., 2004;Jiang et al., 2007;Martignago et al., 2019). Loss-of-function *fld* mutants exhibit a characteristic late-flowering phenotype resulting from elevated expression of the floral repressor *FLC* (He et al., 2003;He et al., 2004;Jiang et al., 2007). The delayed flowering observed in *spm4-1*, *spm4-2*, and *spm4-3* is therefore consistent with previously characterized *fld* mutants and supports the conclusion that these alleles disrupt FLD function. Importantly, previous studies have demonstrated that the immune defects associated with *fld* mutants cannot be explained simply because of altered flowering time. For example, mutations affecting other flowering regulators do not phenocopy the immune phenotypes of *fld*, and restoration of flowering through manipulation of flowering pathways does not restore normal defense responses in *fld* backgrounds (Singh et al., 2013). Thus, the role of FLD in plant immunity appears to be genetically separable from its function in flowering-time regulation. Our finding that loss of FLD abolishes the hyperaccumulation of Pip/NHP in *pmr4* further supports the notion that FLD directly participates in immune regulatory pathways rather than indirectly affecting defense through developmental changes.

A role for FLD in plant immunity was first revealed through its identification as RSI1 (Required for Salicylic Acid Induction 1) in screens for mutants defective in SAR (Singh et al., 2013). The *rsi1/fld* mutant displays compromised SAR and reduced induction of defense-associated genes following pathogen infection. Subsequent studies showed that FLD contributes to systemic immunity, pathogen-induced defense gene expression, and chromatin modifications associated with immune activation (Singh et al., 2014b;Singh et al., 2023;Kumar et al., 2025). However, despite extensive genetic evidence linking FLD to immunity, the underlying molecular mechanisms have remained poorly understood. Several models have been proposed, including direct regulation of defense gene chromatin states, modulation of SA-associated signaling pathways, and broader roles in transcriptional reprogramming during immune activation (Singh et al., 2014b;Singh et al., 2023;Kumar et al., 2025). Nevertheless, a unifying explanation accounting for the diverse immune-related phenotypes of *fld/rsi1* mutants has been lacking.

The discovery that FLD is required for NHP overaccumulation in *pmr4* provides a potential mechanistic framework for understanding many of the previously reported immune functions of FLD. NHP has emerged as a central regulator of both local and systemic immunity and is now recognized as a critical signal for SAR establishment (Hartmann et al., 2018;Hartmann and Zeier, 2019;Yildiz et al., 2021;Zeier, 2021;Shields et al., 2022;Li et al., 2023;Löwe et al., 2023;Yildiz et al., 2023). Biosynthesis of NHP through the ALD1–SARD4–FMO1 pathway is strongly induced during immune activation, and mutants defective in NHP production exhibit profound defects in SAR (Hartmann and Zeier, 2019;Huang et al., 2020;Yildiz et al., 2021). Because *fld/rsi1* mutants also show compromised systemic immunity, it is tempting to speculate that FLD promotes SAR at least in part through regulation of NHP biosynthesis, accumulation, or signaling. In this model, loss of FLD would reduce activation of the NHP pathway, leading to defects in systemic defense amplification and SAR establishment. Such a mechanism would readily explain the previously reported requirement of FLD/RSI1 for systemic immunity and would place FLD within a central regulatory node controlling one of the most important immune signaling pathways in plants.

The biochemical properties of FLD further support a potential epigenetic mechanism for regulation of the NHP pathway. FLD contains a conserved amine oxidase domain and is closely related to lysine-specific histone demethylases, functioning as a chromatin regulator that influences gene expression through modification of histone methylation states (He et al., 2003;Jiang et al., 2007). At the *FLC* locus, FLD promotes transcriptional repression through chromatin remodeling and removal of active histone marks (He et al., 2003;He et al., 2004;Jiang et al., 2007). Numerous studies have now demonstrated that epigenetic regulation plays crucial roles in plant immunity (Jaskiewicz et al., 2011;Luna et al., 2012;Singh et al., 2014a;Lopez Sanchez et al., 2016;Ramirez-Prado et al., 2018;Zhang et al., 2018;Xie and Duan, 2023). Dynamic changes in DNA methylation and histone modifications occur following pathogen infection and contribute to the activation of defense genes. Histone acetylation, methylation, and chromatin remodeling factors have all been implicated in immune priming, defense gene induction, and SAR (Jaskiewicz et al., 2011;Luna et al., 2012;Singh et al., 2014a;Ramirez-Prado et al., 2018;Xie and Duan, 2023). Furthermore, several defense-associated genes exhibit chromatin changes that facilitate rapid transcriptional activation during subsequent pathogen challenge, providing a molecular basis for immune memory and priming (Jaskiewicz et al., 2011;Luna et al., 2012;Ramirez-Prado et al., 2018;Xie and Duan, 2023).

Within this context, FLD may function as an epigenetic regulator required for activation of genes involved in NHP biosynthesis or signaling. One possibility is that FLD directly regulates chromatin states at key NHP pathway genes such as *ALD1*, *SARD4*, *FMO1*, or upstream transcriptional regulators controlling their expression. Alternatively, FLD may influence broader chromatin landscapes that are necessary for immune amplification and systemic signaling. Although our data do not distinguish between these possibilities, the strong reduction of PIP accumulation in the *gsl5-1 fld* double mutants clearly demonstrates that FLD is required for full activation of the NHP pathway (Figure 5). Given the extraordinary increase in PIP observed in *pmr4*, even relatively modest epigenetic effects on NHP biosynthetic gene expression could have profound consequences for metabolite accumulation and downstream defense responses.

In summary, our results identify FLD as an essential positive regulator of NHP accumulation and provide a mechanistic link between epigenetic regulation and immune-metabolite signaling. Together with our previous characterization of *PAD4*, *ALD1*, and *FMO1* as suppressors of *pmr4*, these findings establish a genetic framework in which FLD acts upstream of NHP accumulation to promote the autoimmune phenotype of *pmr4*. More broadly, this work strengthens the emerging view that chromatin-based regulation is an integral component of plant immune signaling networks and suggests that epigenetic regulators such as FLD contribute to growth-defense tradeoffs by controlling the activation of potent immune metabolites including NHP. Future studies aimed at identifying the direct chromatin targets of FLD during immune activation should provide important insights into how epigenetic mechanisms regulate NHP biosynthesis, SAR, and the balance between defense and plant growth.

## MATERIALS AND METHODS

### Plant materials and growth conditions

Arabidopsis (*Arabidopsis thaliana*) WT and mutant plants used in the study are all in the *Col-0* background. Homozygous T-DNA insertion mutant *gsl5-1* (GABI_089H05) has been previously described (Fan et al., 2025). Arabidopsis plants were grown in growth chambers or rooms at 24°C, 70 μmol m^-2^s^-1^ light on a photoperiod of 12-hour light and 12-hour dark as previously described (Fan et al., 2025).

### Chemical mutagenesis, suppressor screens and mapping by BSA-NGS

EMS mutagenesis of *Arabidopsis thaliana gsl5-1* seeds and screening for *suppressor of spm* mutants were performed as previously described (Fan et al., 2025). To identify the causal mutation in *spm4-1*, the *gsl5-1 spm4-1* suppressor mutant was crossed with the parental *gsl5-1* line to generate an F₂ mapping population. Individual F₂ plants were phenotyped at 9-10 weeks after germination and classified based on the presence or absence of the characteristic *gsl5-1* growth defects, including spontaneous lesion formation and premature senescence. Leaf tissues from 50–60 plants of each phenotypic class were collected separately and pooled for genomic DNA (gDNA) extraction.

The pooled gDNA samples were subjected to whole-genome resequencing using the Illumina sequencing platform. Sequence reads were processed for quality control and aligned to the *Arabidopsis thaliana* reference genome (TAIR10). Single-nucleotide polymorphisms (SNPs) and insertion/deletion polymorphisms (indels) were identified and used for bulked segregant analysis. Variant allele frequencies were calculated across the genome to identify chromosomal regions linked to the suppressor mutation, as previously described. Candidate mutations within the linked interval were subsequently examined to identify the causal gene underlying the *spm4-1* phenotype.

### Molecular complementation of the *gsl5 spm4* suppressor mutant

A WT genomic fragment containing the entire *FLD* coding sequence was amplified by PCR from WT genomic DNA using *FLD*-specific primers (5’-agcgtcgacggccttgttgaaaccaccatg-3’ and 5’-agcttaattaagagctgatgatttggcggag-3’). The fragment was digested with SalI and PacI and cloned into the XhoI and PacI sites behind the *CaMV 35S* promoter in the binary vector pFGC5941 (Kerschen et al., 2004). The resulting construct was introduced into *Agrobacterium tumefaciens* strain GV3101 and transformed into *gsl5-1 spm4-1* plants using the floral-dip method (Bent, 2006). Transgenic plants were selected for Basta resistance and subsequently transferred to soil for phenotypic analysis. Independent T₁ transformants were analyzed for *FLD* transcript levels by real-time quantitative PCR (RT-qPCR) and evaluated for restoration of the *gsl5-1* autoimmune phenotype, including reduced growth, spontaneous necrosis, and premature senescence.

### RT-qPCR

Total RNA isolation, DNase treatment, cDNA synthesis and RT-qPCR using *FLD*-specific primers (5’-ggtctcattctccgcaccaa-3’ and 5’-gacctacgttagggacggga-3’) were performed with *ACTIN2* (5’-accagctcttccatcgagaa-3’ and 5’-gggcatctgaatctctcagc-3’) as an internal control as described previously (Fu et al., 2023).

### Generation of additional *spm4/fld* mutant alleles using CRISPR/cas9

To generate independent loss-of-function alleles of *FLD*, two guide RNAs (gRNAs) (5’-tctccgcaccaaagaaacga-3’ and 5’-ctttgcttgctctcactgct-3’) targeting the first exon near the 5′ end of the *FLD* coding sequence were designed using the CRISPR-P online tool (Liu et al., 2017). The gRNA sequences were assembled into a plant CRISPR/Cas9 expression vector with the cas9 gene driven by an egg cell-specific promoter (Zheng et al., 2020). The CRISPR/Cas9 construct was introduced into *A. tumefaciens* strain GV3101. The resulting construct was transformed into *gsl5-1* plants using the floral-dip method (Bent, 2006). Transgenic T₁ plants were selected for Basta resistance and screened for mutations at the target sites by PCR amplification and DNA sequencing. Two independent edited lines carrying single-nucleotide deletion or insertions at nucleotide position 184 downstream of the *FLD* start codon were identified. These mutant alleles were designated *spm4-2* and *spm4-3*, respectively. Homozygous mutant lines were identified by sequencing and analyzed for suppression of the *gsl5-1* growth-defect phenotype.

### Quantification of PIP

Preparation of leaf tissues, extraction and quantification of PIP by high-performance liquid chromatography-tandem MS (HPLC-MS/MS) were performed as previously described (Semeraro et al., 2015;Fan et al., 2025).

## ACKNOWLEDGMENTS

We thank the Arabidopsis Resource Center at the Ohio State University for providing the mutant used in the study. We thank other members of the Chen lab for help in genotyping of T-DNA mutants, analysis of PIP and isolation of RNA. This work was supported by US National Science Foundation (grants no. IOS1758767 and MCB2241515).

## AUTHOR CONTRIBUTIONS

Z.C. conceived the original research plans. B.F. performed most of the experiments; Z.C. performed some of the experiments; B.F. and Z.C. wrote the article.

